# ESR2 orchestrates cytokinin dynamics leading to developmental reprogramming and green callus formation

**DOI:** 10.1101/2024.06.20.596278

**Authors:** Yolanda Durán-Medina, David Díaz-Ramírez, Humberto Herrera-Ubaldo, Maurizio Di Marzo, Andrea Gómez Felipe, J. Erik Cruz-Valderrama, Carlos A. Vázquez, Herenia Guerrero-Largo, Lucia Colombo, Ondrej Novak, Stefan de Folter, Nayelli Marsch-Martínez

## Abstract

Callus formation and shoot regeneration are natural plant abilities triggered by stress and damage. They are also key components of tissue culture, which for many species is crucial for gene editing, transformation, propagation, and other technologies, and their study provides valuable insights into plant development. The transcription factor ENHANCER OF SHOOT REGENERATION 2 (ESR2/DRNL/BOL/SOB) promotes green callus formation in roots and shoot regeneration when overactive, while the phythormone cytokinin plays a prominent role in both phenomena. Yet, the positive action of ESR2 on the cytokinin pathway had not been previously described. We explored the effects of ESR2 and found that cytokinin content and the expression of the cytokinin biosynthesis gene *ISOPENTENYLTRANSFERASE 5* (*IPT5)* increase in plants where ESR2 activity is induced. ESR2 also regulates the cytokinin signaling repressor *ARABIDOPSIS HISTIDINE PHOSPHOTRANSFER 6* (*AHP6),* and surprisingly, the ESR2-stimulated green calli formation requires both *IPT5* and *AHP6*. Therefore, ESR2 promotes both cytokinin biosynthesis and cytokinin signaling inhibition and requires a paradoxical combination of these two for green callus induction. This finding provides a foundation to better understand the processes involved in tissue reprogramming towards callus formation and the role of ESR2 in shoot regeneration and the development of new organs.

## Introduction

The remarkable regenerative capacity of plants allows the emergence of new organs or complete plants from a small piece of somatic tissue. This capacity highlights the incredible plasticity of plant cells, which has long attracted the interest of researchers (Ibáñez et al., 2020; Ikeuchi et al., 2016). Great advance in the study of plant cellular plasticity occurred after the discovery of the phytohormones auxin and cytokinin and their application to *in vitro* plant tissue culture (Melnyk, 2023). The auxin/cytokinin phytohormone ratio is key for the fate specification of plant cells. In 1957, Skoog and Miller’s pioneering work using tobacco explants revealed that high ratios of auxin-to-cytokinin or cytokinin-to-auxin induce root or shoot regeneration, respectively, while an intermediate ratio of auxin and cytokinin promotes a mass of growing cells known as “callus” (Skoog & Miller, 1957). Since then, callus has been widely used as a valuable tool for the propagation and genetic engineering of a wide variety of plants (Ikeuchi et al., 2013). This type of tissue benefits from indefinite growth while it is maintained in a callus-inducing medium, but when it is moved to a new medium with the right combination of cytokinin and auxin, it is possible to obtain shoot or root regeneration (Skoog & Miller, 1957). Recent research has advanced the understanding of the molecular mechanisms underlying cytokinin’s function in the control of green callus formation and shoot regeneration (reviewed in Šmeringai et al., 2023).

The AP2/ERF2 ENHANCER OF SHOOT REGENERATION (ESR) transcription factors can also trigger callus and shoot formation. The overexpression of ESR1/DORNRÖSCHEN (DRN) in Arabidopsis promotes cytokinin-independent shoot generation and increases shoot regeneration efficiency in the presence of cytokinin (Banno et al., 2001). ESR2/BOLITA (BOL) overexpression or gain of function also induces callus and shoot production without exogenous phytohormone application, and it is more active than ESR1 in promoting shoot regeneration in tissue culture (Ikeda et al., 2006; Marsch-Martinez et al., 2006). ESR2 is also known as DORNRÖSCHEN-LIKE (DRNL) (Chandler & Werr, 2014; Kirch et al., 2003) and SUPPRESSOR OF PHYTOCHROMEB-4 2 (SOB2) (Ward et al., 2006). In this work, focused on callus development, it will be referred to as ESR2. The *ESR2* expression pattern precedes and partially overlaps local auxin maxima, as revealed by the synthetic auxin response reporter DR5 (Chandler et al., 2011; Chandler & Werr, 2014), and ESR2 indirectly regulates auxin biosynthesis through the transcription factor STYLISH1 (STY1) (Eklund et al., 2011). However, the effects of increased ESR2 expression closely resemble the effects of increased cytokinin levels, such as the dark green color in leaves (Cortleven & Schmülling, 2015), primary root growth inhibition (Dello Ioio et al., 2007; Liu et al., 2022; Werner et al., 2003), and green callus development (Ikeuchi et al., 2013; Sakai et al., 2001). The ESR2 induction phenotype has been described as comparable to the phenotype of mutants overproducing cytokinins (Ikeda et al., 2006). ESR2 overexpression also affects the expression of genes related to the cytokinin pathway (Ikeda et al., 2006a; Marsch-Martinez et al., 2006). Our previous research on gynoecium development indicated a positive regulatory effect of ESR2 on the cytokinin pathway at the early stages of gynoecium development (Durán-Medina et al., 2017a). Furthermore, the mutation of the tomato ortholog of *ESR2*, *LEAFLESS (LFS),* alters the expression of cytokinin biosynthesis and signaling genes, suggesting a positive regulatory role (Capua & Eshed, 2017). Therefore, based on this data, this study aims to investigate the role of ESR2 as a positive regulator of the cytokinin pathway (biosynthesis or signaling) leading to green callus formation.

## RESULTS

### ESR2 induction promotes severe defects in seedling organ development

One of the most striking reported effects of *ESR2* is the formation of green calli in roots of young seedlings where the gene is constitutively overexpressed, or in root explants where the overexpression was induced (Ikeda et al., 2006a; Marsch-Martinez et al., 2006). To enhance our understanding about the effects of this transcription factor in plant development, we analyzed the morphological changes in whole seedlings caused by the induction of ESR2 activity after germination in the *35S::ESR2:ER* (*ESR2:ER*) inducible line. In this line, *β*-estradiol triggers the entrance of the ESR2 transcription factor into the nucleus, where it regulates gene expression (Eklund et al., 2011; Ikeda et al., 2006). As expected, the induction of ESR2 activity resulted in altered development of aerial and root tissues (Fig. 1A-H). The severity of these morphological alterations depended on the plant developmental stage in which this transcription factor was activated, and the alterations became more evident with the increase in induction time. The induction of ESR2 in seedlings led to an arrest of leaf development (Fig. 1A-D). The induced seedlings did not produce new leaves like those of Wild Type (WT) seedlings, and the leaves that were in early developmental stages at the beginning of the treatment did not expand as non-induced leaves. Moreover, they acquired a dark green color in induced seedlings compared to non-induced ones (Fig. 1A-D). In roots, the developmental changes were also profound. The root apex thickened, coiled, and began to acquire a green color (Fig. 1G-H). The growth of the primary root was arrested, and the induced seedlings had shorter primary roots and a higher number of thick “lateral roots” than non-induced seedlings (Fig 1E). These new thickened lateral roots displayed stunted growth. The induced seedlings had a root length of 37 mm and 8 lateral roots on average, per seedling, while the non-induced seedlings had a root length of 76 mm and 5 lateral roots on average (Fig. 1I-J). In addition, we also observed disorganized lateral root positioning along the main root axis, with some roots growing at an unusual proximity to each other (Fig. 1H). At longer periods of ESR2 induction (21 days), the entire root demonstrated green callus formation. These green calli developed initially at some distance from each other, at the tip or the base of lateral roots, and subsequently, the whole root became green (Fig. 1F).

**Figure 1.**
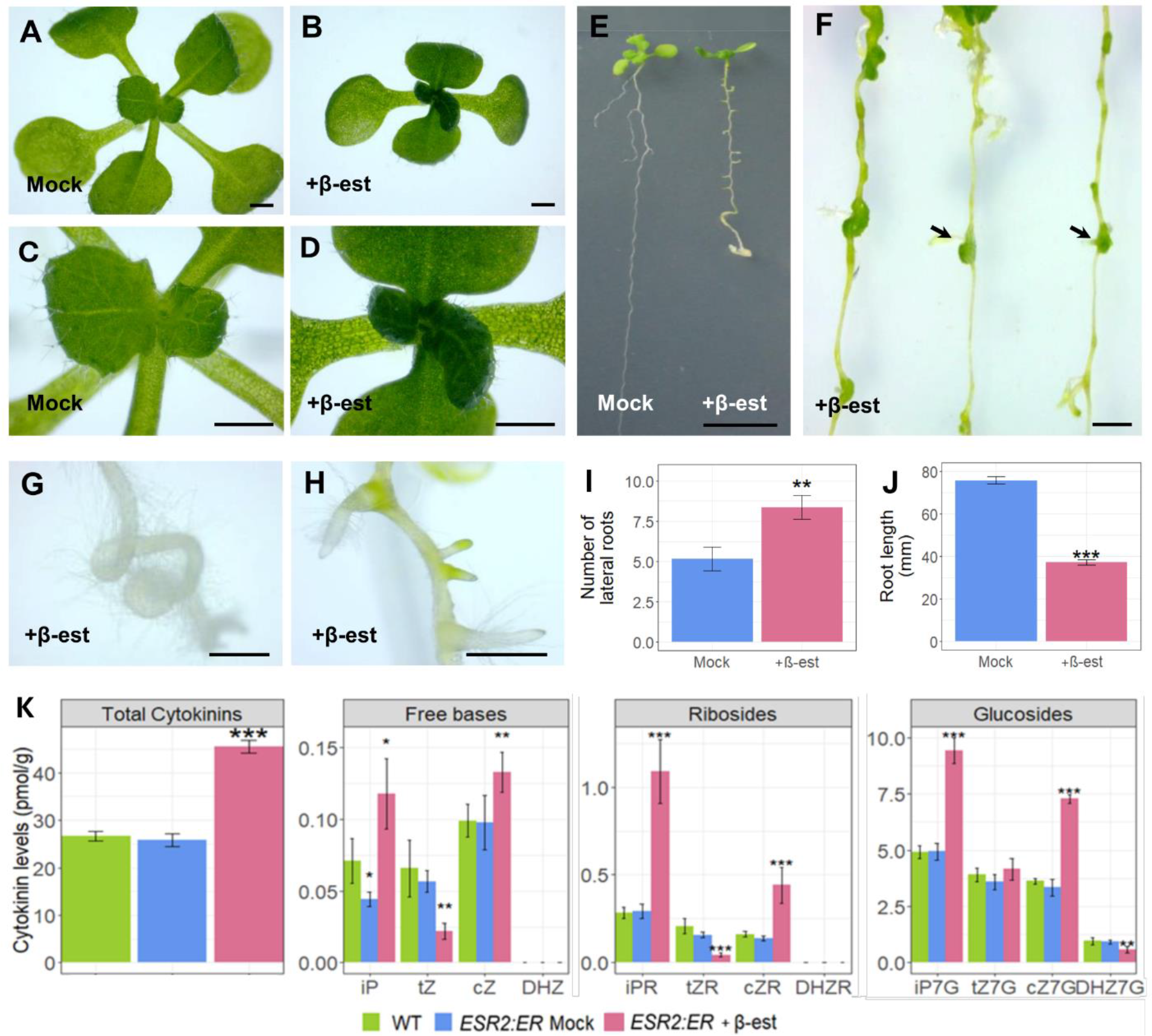
Effects of ESR2 overactivity in Arabidopsis seedling development. A) Aerial tissue of mock and B) induced seedlings 15 days after ESR2 induction. C) Closeup of the younger leaves in mock and D) induced seedlings 15 days after ESR2 induction. E) Comparison between roots of induced and mock treated seedlings. F) Calli developing from the tissue near or at the base of young lateral roots after 21 days of ESR2 induction. G) Closeup of the main root apex of induced seedlings. H) Disorganization of lateral root emergence in induced seedlings. I) Average number of lateral roots. Mann-Whitney test, **p ≤ 0.001. J) Average of primary root length. Student’s t- test, *** p ≤ 0.0001. K) Cytokinin levels in WT and ESR2 induced and not induced seedlings. Student’s t-test, *, **, and ***, p ≤ 0.01, p ≤ 0.001, and p ≤ 0.001, respectively. Scale bar (A-D, G, H) 500 um, (E) 1 cm and (F) 1 mm.

In summary, the induction of ESR2 in whole seedlings affected the development of leaves and roots, reducing or inhibiting their growth and leading to green callus development first in regions associated with lateral roots and later along the whole root.

### Cytokinin endogenous levels increase after ESR2 induction in seedlings

The effects of ESR2 induction were similar to those caused by increased levels of cytokinins, and to ascertain the presence of an increased content of these phytohormones in the induced plants, we conducted a quantitative analysis of endogenous cytokinin levels in WT versus induced and non-induced *ESR2:ER* seedlings (Fig. 1K). The quantification revealed a significant increase in total endogenous cytokinin levels (45.46 pmol/g FW ± 1.38) in *ESR2:ER* induced seedlings in comparison to *ESR2:ER* non-induced seedlings (25.84 ± 1.39) and wt seedlings (26.67 ± 1.02) (Table 1), which had a similar total cytokinin content. The cytokinin types that showed statistically significant accumulation in induced *ESR2:ER* seedlings were isopentenyl adenine (iP) and cis-zeatin (cZ) types (free bases, ribosides, and glucosides), whereas the level of some trans-zeatin (tZ) type cytokinins (free bases and ribosides) had actually decreased (Fig. 1K, Table 1). Therefore, the induction of ESR2 leads to an increase in total cytokinin levels, particularly isopentenyl adenine and cis-zeatin species, along with a reduction of trans-zeatin free bases and ribosides.

**Table 1.** CK quantification.

| Genotype | Total Cytokinins | SE |  | CK Bases | SE |  | CK Ribosides | SE |  | CK Nucleotides | SE |  | CK O-glucosides | SE |  | CK N-glucosides | SE |  |
| --- | --- | --- | --- | --- | --- | --- | --- | --- | --- | --- | --- | --- | --- | --- | --- | --- | --- | --- |
| Col-0 | 26.667 | ±1.018 |  | 0.236 | ±0.021 |  | 0.652 | ±0.078 |  | 5.773 | ±0.789 |  | 2.575 | ±0.145 |  | 17.431 | ±0.400 |  |
| ESR2 MOCK | 25.845 | ±1.386 |  | 0.199 | ±0.023 | * | 0.588 | ±0.047 |  | 5.800 | ±0.589 |  | 2.461 | ±0.211 |  | 16.798 | ±1.233 |  |
| ESR2 Induced | 45.461 | ±1.379 | *** | 0.273 | ±0.032 |  | 1.573 | ±0.282 | *** | 13.180 | ±1.017 | *** | 4.495 | ±0.386 | *** | 25.940 | ±1.648 | *** |

| Genotype | Total iP-types | SE |  | iP | SE |  | iPR | SE |  | iPRMP | SE |  | iP7G | SE |  | iP9G | SE |  |
| --- | --- | --- | --- | --- | --- | --- | --- | --- | --- | --- | --- | --- | --- | --- | --- | --- | --- | --- |
| Col-0 | 8.244 | ±0.248 |  | 0.071 | ±0.016 |  | 0.283 | ±0.030 |  | 2.179 | ±0.212 |  | 4.919 | ±0.294 |  | 0.792 | ±0.028 |  |
| ESR2<br>MOCK | 8.570 | ±0.878 |  | 0.044 | ±0.005 | * | 0.291 | ±0.041 |  | 2.516 | ±0.585 |  | 4.935 | ±0.371 |  | 0.784 | ±0.038 |  |
| ESR2<br>Induced | 18.345 | ±0.670 | *** | 0.118 | ±0.024 | * | 1.087 | ±0.182 | *** | 6.198 | ±0.525 | *** | 9.408 | ±0.569 | *** | 1.534 | ±0.089 | *** |

| Genotype | Total tZ-types | SE |  | tZ | SE |  | tZR | SE |  | tZRMP | SE |  | tZOG | SE |  | tZROG | SE |  | tZ7G | SE |  | tZ9G | SE |
| --- | --- | --- | --- | --- | --- | --- | --- | --- | --- | --- | --- | --- | --- | --- | --- | --- | --- | --- | --- | --- | --- | --- | --- |
| Col-0 | 9.732 | ±0.556 | | 0.066 | ±0.020 | | 0.207 | ±0.043 | | 0.894 | ±0.199 | | 1.385 | ±0.093 | | 0.110 | ±0.018 | | 3.902 | ±0.305 | | 3.168 | ±0.268 |
| ESR2 MOCK | 9.043 | ±0.632 | | 0.057 | ±0.008 | | 0.158 | ±0.015 | | 0.752 | ±0.159 | | 1.232 | ±0.160 | | 0.100 | ±0.005 | | 3.581 | ±0.334 | | 3.163 | ±0.349 |
| ESR2 Induced | 8.026 | ±1.230 | * | 0.022 | ±0.006 | ** | 0.044 | ±0.010 | *** | <LOD | | | 0.690 | ±0.082 | *** | 0.171 | ±0.015 | *** | 4.159 | ±0.486 | | 2.939 | ±0.665 |

| Genotype | Total cZ-types | SE |  | cZ | SE |  | cZR | SE |  | cZRMP | SE |  | cZOG | SE |  | cZROG | SE |  | cZ7G | SE |  | cZ9G | SE |
| --- | --- | --- | --- | --- | --- | --- | --- | --- | --- | --- | --- | --- | --- | --- | --- | --- | --- | --- | --- | --- | --- | --- | --- |
| Col-0 | 7.663 | ±0.398 | | 0.099 | ±0.011 | | 0.162 | ±0.015 | | 2.699 | ±0.420 | | 0.247 | ±0.016 | | 0.833 | ±0.047 | | 3.623 | ±0.114 | | <LOD | |
| ESR2 MOCK | 7.241 | ±0.587 | | 0.098 | ±0.019 | | 0.139 | ±0.015 | | 2.531 | ±0.207 | | 0.250 | ±0.027 | | 0.879 | ±0.049 | | 3.344 | ±0.368 | | <LOD | |
| ESR2 Induced | 18.473 | ±0.815 | *** | 0.133 | ±0.014 | ** | 0.441 | ±0.101 | *** | 6.982 | ±0.619 | *** | 0.440 | ±0.016 | *** | 3.194 | ±0.457 | *** | 7.284 | ±0.201 | *** | <LOD | |

| Genotype | Total<br>DHZ-types | SE |  | DHZ | SE |  | DHZR | SE |  | DHZRMP | SE |  | DHZOG | SE |  | DHZROG | SE |  | DHZ7G | SE |  | DHZ9G | SE |  |
| --- | --- | --- | --- | --- | --- | --- | --- | --- | --- | --- | --- | --- | --- | --- | --- | --- | --- | --- | --- | --- | --- | --- | --- | --- |
| Col-0 | 1.027 | ±0.161 |  | <LOD |  |  | <LOD |  |  | <LOD |  |  | <LOD |  |  | <LOD |  |  | 0.962 | ±0.152 |  | 0.065 | ±0.010 |  |
| ESR2<br>MOCK | 0.992 | ±0.084 |  | <LOD |  |  | <LOD |  |  | <LOD |  |  | <LOD |  |  | <LOD |  |  | 0.921 | ±0.085 |  | 0.071 | ±0.008 |  |
| ESR2<br>Induced | 0.616 | ±0.142 | ** | <LOD |  |  | <LOD |  |  | <LOD |  |  | <LOD |  |  | <LOD |  |  | 0.572 | ±0.136 | ** | 0.044 | ±0.009 | * |

### *IPT5* expression increases after *ESR2* activation

Since ESR2 promotes an increase in total cytokinin endogenous levels, we investigated whether ESR2 could regulate genes involved in cytokinin biosynthesis. The first and rate-limiting step in cytokinin biosynthesis is catalyzed by Isopentenyl Transferases (IPTs) (Miyawaki et al., 2004, 2006). *IPT5* expression has been reported at the early stage I of lateral root development and *de novo* root development (Bustillo-Avendaño et al., 2018; Chang et al., 2015; Miyawaki et al., 2004) and during the process of callus development from roots (Atta et al., 2009). Based on this, we employed RT-qPCR to assess whether ESR2 overactivation affected *IPT5* expression. An increase in *IPT5* expression was observed while comparing WT and *ESR2:ER* seedlings treated with β- estradiol and cycloheximide (which inhibits translation) or a mock solution for 30 min at 9 Days After Germination (DAG) (Fig. S1A). To follow the expression at later times after induction, and visualize its spatial pattern, we analyzed the *IPT5* promoter marker line *IPT5::GUS* (Dello Ioio et al., 2008) in the *ESR2:ER* background. A clear increase in GUS signal in the vascular region of hypocotyls, cotyledons, and leaves, was observed 48 hours after ESR2 induction (Fig. 2A). Interestingly, in non-induced *ESR2:ER IPT5::GUS* seedlings, the GUS signal was slightly higher in some vascular regions when compared to the WT background (Fig. S1B), suggesting a slight leakage of ESR2:ER activity. Nevertheless, after ESR2 induction, GUS staining intensified and expanded completely throughout the entire vasculature of the aerial organs (Fig. 2A). The induction of ESR2 activity also altered *IPT5* expression in roots. In non-induced roots, it was detected in lateral root primordia and was maintained at their apex as the young lateral root elongated (Fig. 2D and S2, Miyawaki et al., 2004), while the apex of the main root did not show *IPT5* expression. However, when ESR2 was activated, *IPT5* expression was observed in the main root apex (Fig. 2A and 2D). At longer induction times, increased *IPT5* expression was detected in the vascular region, and six to eight days after induction, the expression had extended along the whole root as two continuous lines in the vasculature region (Fig. 2B and D). In conclusion, ESR2 activation leads to a significant increase in *IPT5* expression.

**Figure 2.**
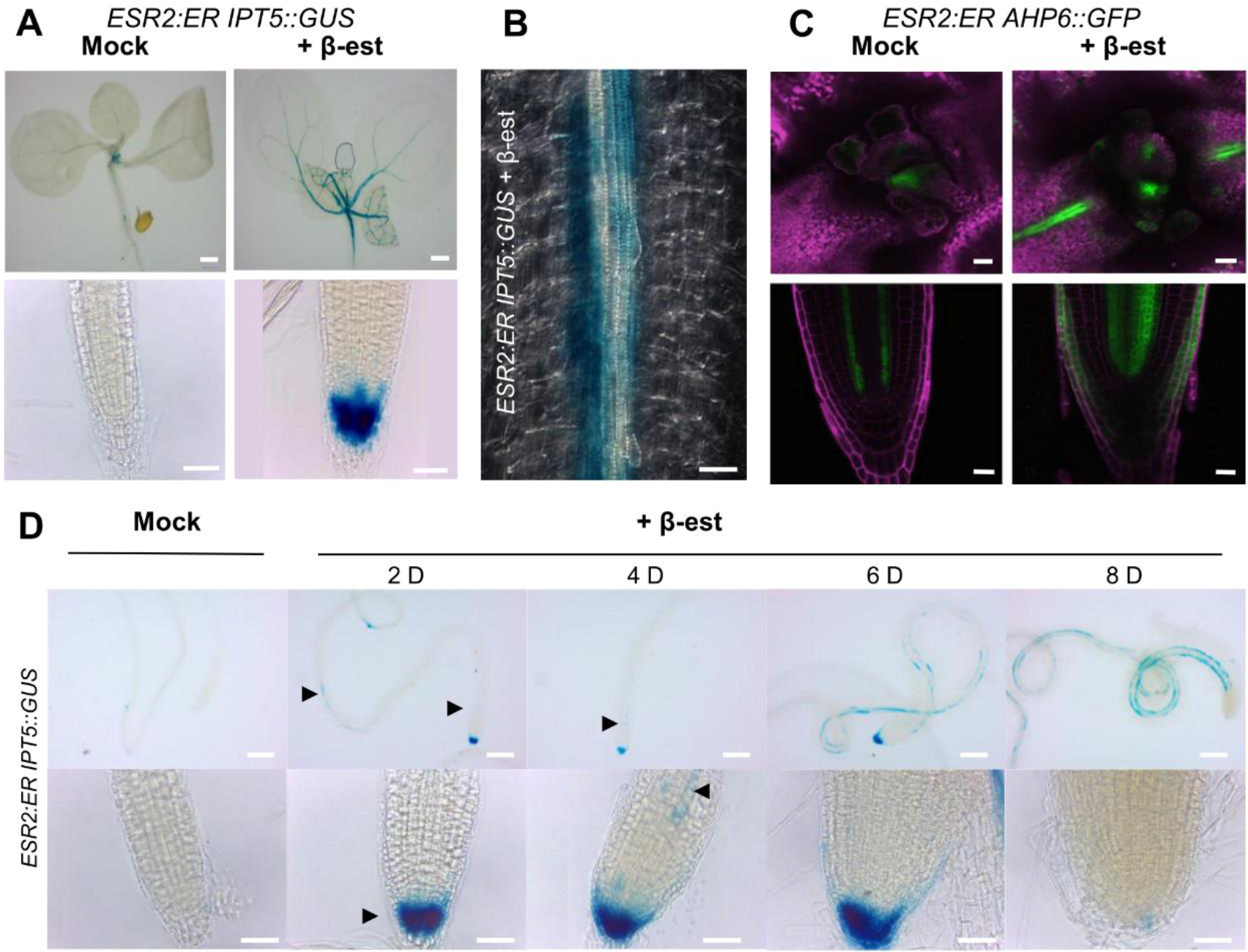
*IPT5* and *AHP6* regulation by ESR2 in aerial and root vascular tissues. A) *IPT5* expression in aerial tissue and the root tip of mock treated and induced ESR2::ER, 48 hours after treatment. **B)** *IPT5* expression in the vasculature of the ESR2::ER main root after 8 days of ESR2 induction. **C)** *AHP6* expression in aerial tissue and the root tip 48 hours after ESR2 induction compared to mock-treated seedlings. **D)** Sequential changes in *IPT5* expression in the main root from 2 to 8 days after ESR2 induction. IPT5 is not expressed at the root apex in non-induced seedlings, but it can be detected after ESR2 induction for a few days, and then it disappears again. It also expands in the vascular region along the root with time. Arrows indicate first signs of *IPT5* ectopic expression in the root after ESR2 induction. Scale bars A) 0.5 mm in aerial tissue, 0.05 mm in roots; B) 0.02 mm; D) 0.2 mm in above pictures and 0.05 mm in below pictures.

### *AHP6* expression increases after ESR2 induction

ESR2 activation also promotes the ectopic expression of *AHP6*, involved in cytokinin signaling, in the gynoecium pro-vasculature (Durán-Medina et al., 2017a), and increased expression in root explants (Ikeda et al., 2006). Therefore, given its relation to the cytokinin pathway, we analyzed *AHP6* expression after ESR2 induction in seedlings through RT-qPCR and found that it increased 30 minutes after activation of ESR2 (Fig. S1A).

While analyzing the *AHP6* promoter reporter line *AHP6::GFP* (Mähönen et al., 2006) in the *ESR2:ER* background, we observed a clear increase of *AHP6* expression associated with vascular tissues of petioles and roots 48 hours after ESR2 induction, similar to *IPT5* (Fig. 2C). Interestingly, 48 hours after ESR2 induction, a higher number of vascular cells with *AHP6* expression were observed in roots. *AHP6* also presented ectopic expression in epidermal cells at the root apex (Fig. 2C).

Taken together, the results indicate that ESR2 activation caused increased and ectopic expression of *IPT5* and *AHP6* in young seedlings.

### The loss of *IPT5* and *AHP6* functions suppress morphological defects caused by ESR2 overactivity in a gene-dependent manner

After observing the increase in *IPT5* and *AHP6* expression due to ESR2 induction, we assessed their involvement in the phenotypes caused by ESR2 induction. For this, we analyzed how the loss of *IPT5* or *AHP6* function affected the phenotypes triggered by ESR2 activation in *ESR2:ER ipt5* or *ahp6* mutants. When ESR2 was induced at 6 DAG, seedlings presented stunted cotyledons and two true leaves that were barely visible (Fig. 3A). This effect in young leaves has also been observed in constitutive ESR2 overexpression lines (Marsch-Martinez et al., 2006). While the first two true leaves were only slightly visible in *ESR2*:*ER* induced seedlings, they appeared to be larger in the mutant backgrounds (*ESR2:ER ipt5* and *ESR2:ER ahp6)* (Fig. 3A). *ESR2:ER ipt5* leaves exhibited the fastest development and were the largest among the three genotypes, resembling the leaves of non-induced plants (Fig. 3A). 17 days after germination, induced *ESR2:ER* seedlings had not formed more visible leaves in addition to the first two that remained small. In contrast, *ESR2:ER ipt5* and *ESR2:ER ahp6* seedlings had 4 new leaves on average, resembling the phenotype of non-induced seedlings (Fig. 3A). This indicated that the leaf phenotypes produced by ESR2 activation require functional *IPT5* and *AHP6*.

**Figure 3.**
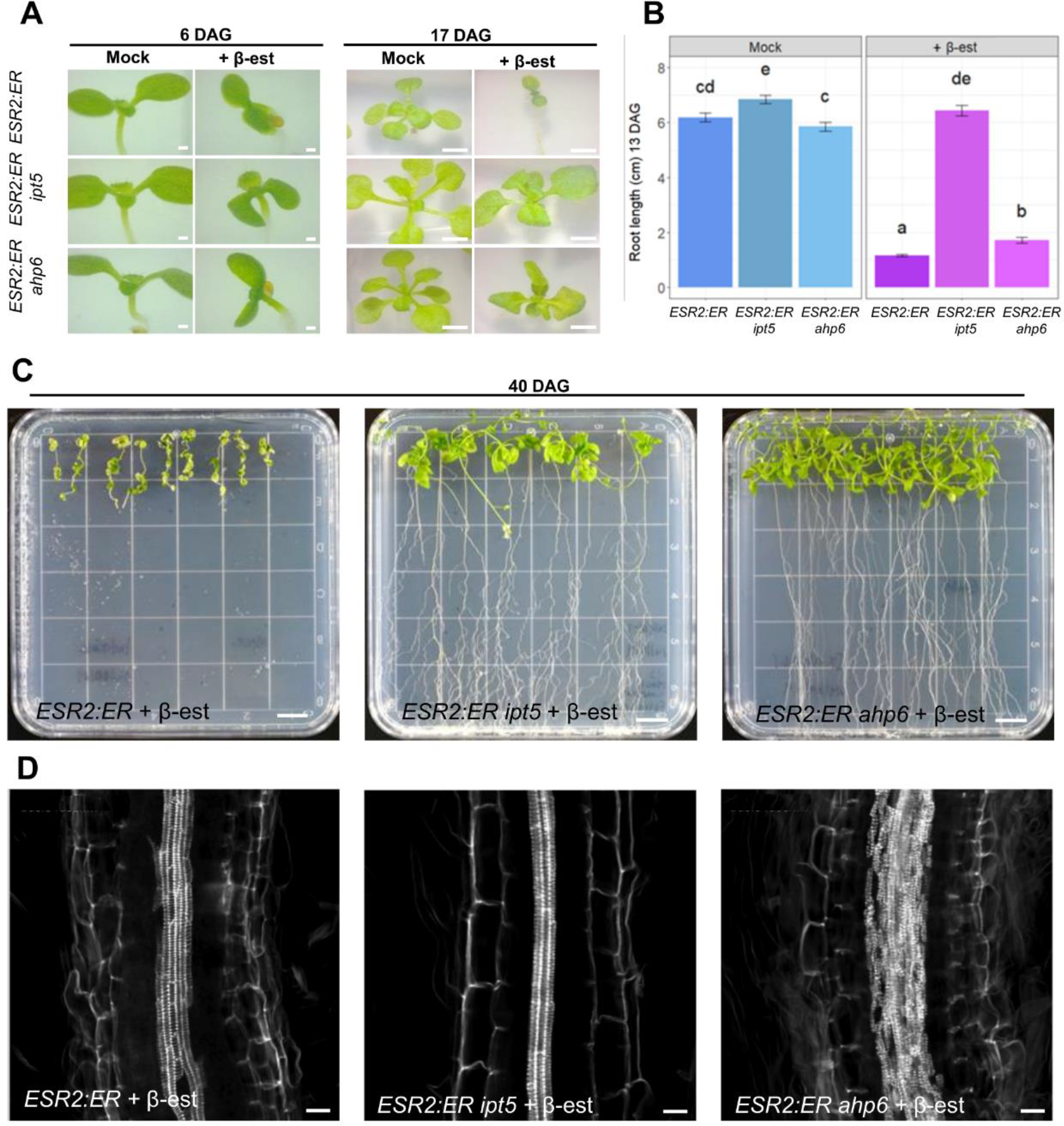
Effects of *IPT5* and *AHP6* loss of function in the ESR2 activation phenotype. A) Aerial tissue development in wild type, *ahp6* and *ipt5* mutant backgrounds at 6 and 17 days after ESR2 induction (+ β-est), compared to not induced seedlings. **B**) Root length of *ESR2:ER, ESR2:ER ipt5* and *ESR2:ER ahp6* seedlings 13 days after direct germination either in mock or ESR2 induction medium. Different letters indicate different statistical groups. C) Development of induced *ESR2:ER*, *ESR2:ER ip5* and *ESR2:ER ahp6* plants 40 DAG. **D)** Vascular tissue in primary roots of induced *ESR2:ER*, *ESR2:ER ipt5* and *ESR2:ER ahp6* seedlings 15 days after induction, stained with basic fuchsin. Scale bars A) 6 DAG seedlings 1mm, C) 1 cm, D) 0.02 mm.

ESR2 overactivity also affects root development. Eight days after germination in the induction medium, the main roots of *ESR2:ER* and *ESR2:ER ahp6* seedlings displayed a profusion of root hairs, while fewer root hairs were observed in *ESR2:ER ipt5* seedlings (Fig. S3A). At 13 DAG in the induction medium, the main root apex of *ESR2:ER* seedlings started to coil (Fig. S3B), and the hairy roots were still visible. Main roots had an average length of 1.2 cm, while non-induced roots were 6 cm long on average (Fig. 3B). The roots of induced *ESR2:ER ahp6* seedlings still showed the profusion of root hairs (Fig. S3B) but were about 1.7 cm long on average, slightly longer in comparison to induced *ESR2:ER* roots (Fig. 3B). Interestingly, the loss of *IPT5* function in induced *ESR2:ER* seedlings allowed the development of the root comparable to non-induced seedlings lacking the profusion of root hairs (Fig. S3B) and exhibiting an average length of 6.4 cm (Fig. 3B). Therefore, the early root growth inhibition and increased root hair density promoted by ESR2 activation appear to require a functional *IPT5*, while *AHP6* seems to play a contributory role, but to a lesser extent, in young plants.

We next explored the effect of the loss of *IPT5* or *AHP6* function on ESR2-promoted callus development. To allow evident callus development, plants were analyzed at 40 DAG in the induction medium. The roots of induced *ESR2:ER* plants did not elongate further, and green calli developed along their entire root length (Fig. 3C). *ESR2:ER ipt5* roots were less affected by ESR2 induction as they continued developing, and became longer than *ESR2:ER* roots (Fig. 3C). This was already evident a few days after induction (Fig. S3B). Furthermore, *ESR2:ER ipt5* roots exhibited a marked reduction in green callus formation after induction (Fig. 3C), with the majority of plants failing to show any callus development in the entire root. In some plants, only a few small calli were formed (Fig. S3C), contrasting to the massive callus formation in *ESR2:ER* roots in the WT background. Moreover, the few calli formed in *ESR2:ER ipt5* roots developed only at the base of a few lateral roots, rather than along the entire root (Fig S3D).

In the case of *ESR2:ER ahp6,* though young roots initially presented an ESR2 induction phenotype that resembled the phenotypes of induced *ESR2:ER* seedlings in a wild-type background, the altered phenotypical characteristics were lost as they developed. *ESR2:ER ahp6* roots developed normally afterward despite the prolonged exposure to the induction medium. Their roots were longer than *ESR2:ER* roots (Fig. 3C). Remarkably, these *ESR2:ER ahp6* plants did not form any visible calli after ESR2 induction, not even after the extended induction period (Fig. 3C). The severe reduction or lack of callus formation in *ipt5* and *ahp6* mutants indicates that these effects caused by ESR2 induction require functional *IPT5* and *AHP6*.

When analyzing the roots of induced *ESR2:ER* seedlings in more detail, we noticed anomalies in the vascular tissues, already evident in the *IPT5* and *AHP6* reporter lines within the induced *ESR2:ER* background (Fig. 2B and C). To further investigate these anomalies, we stained the roots with basic fuchsin, which stains lignin and allows the visualization of xylem cells (Mähönen et al., 2000). In non-induced seedlings, two well-defined stained lines corresponding to protoxylem cells were observed, while a stained broad band was observed in the roots of induced seedlings (Fig. 3D). This suggested that ESR2 activation led to an increase in the number of vascular cells. Interestingly, the roots of induced *ESR2:ER ipt5* seedlings presented a reduction in xylem strands compared to induced *ESR2:ER* seedlings, with their xylem appearing to be more similar to those of non-induced plants. Conversely, the roots of induced *ESR2:ER ahp6* seedlings presented an even more disorganized pattern of protoxylem strands than induced *ESR2:ER* roots (Fig. 3D). This indicates that *IPT5* contributes to the altered vasculature phenotype caused by ESR2 induction, while *AHP6* appears to counteract it.

In summary, functional *IPT5* and *AHP6* are required for the ESR2-induced stunted and darker leaf, late root inhibition, and callus formation phenotypes, while *IPT5* is required for the ESR2-induced altered vascular phenotype and *AHP6* counteracts it.

### Cytokinin application partially restores green callus development in *ESR2:ER ipt5* roots

The previous experiments indicated that ESR2 promotes the development of green calli through the regulation of biosynthesis and negative signaling steps of the cytokinin pathway, suggesting that cytokinin plays a prominent role in ESR2-induced callus formation. Therefore, we investigated the effect of exogenous cytokinin on the inducible *ESR2:ER ipt5* and *ahp6* mutant backgrounds. We specifically evaluated whether exogenous cytokinin could compensate for the loss of *IPT5* function, by complementing the lack of cytokinin biosynthesis and restoring callus formation in the *ESR2:ER ipt5* background. We also explored the effect of exogenous cytokinin in the *ESR2:ER ahp6*, where negative signaling is compromised.

For this, seedlings were grown in media supplemented with 20 µM β-estradiol and transplanted to media with 30 µM 6-benzylaminopurine (BAP) and 20 µM β-estradiol at 6 DAG. Interestingly, after 34 days, all *ESR2:ER ipt5* plants presented calli formation in their hypocotyls, near the root-hypocotyl junction, and at the root apex (Fig. 4A and S4B). The calli development along the root was minimal, not at the extent as observed in induced *ESR2:ER* plants, but it was evident that BAP treatments increased calli formation in these plants, in contrast with plants grown without BAP (Fig. S4A-B). In the case of *ESR2:ER ahp6* plants, most did not present visible calli, and only a few developed green root tips resembling calli (about 20%, Fig. S4B-C). Callus development occurred only at the root apex but not in other regions, like in *ESR2:ER* or BAP-treated *ESR2:ER ipt5* plants. Remarkably, instead of calli, some shoot-like structures were observed at the base of *ESR2:ER ahp6* hypocotyls (Fig. 4A). This peculiar effect had not been observed before in any of the previous experiments and was only found in induced *ESR2:ER ahp6* plants treated with cytokinins, suggesting that in these conditions, *AHP6* suppresses shoot formation.

**Figure 4.**
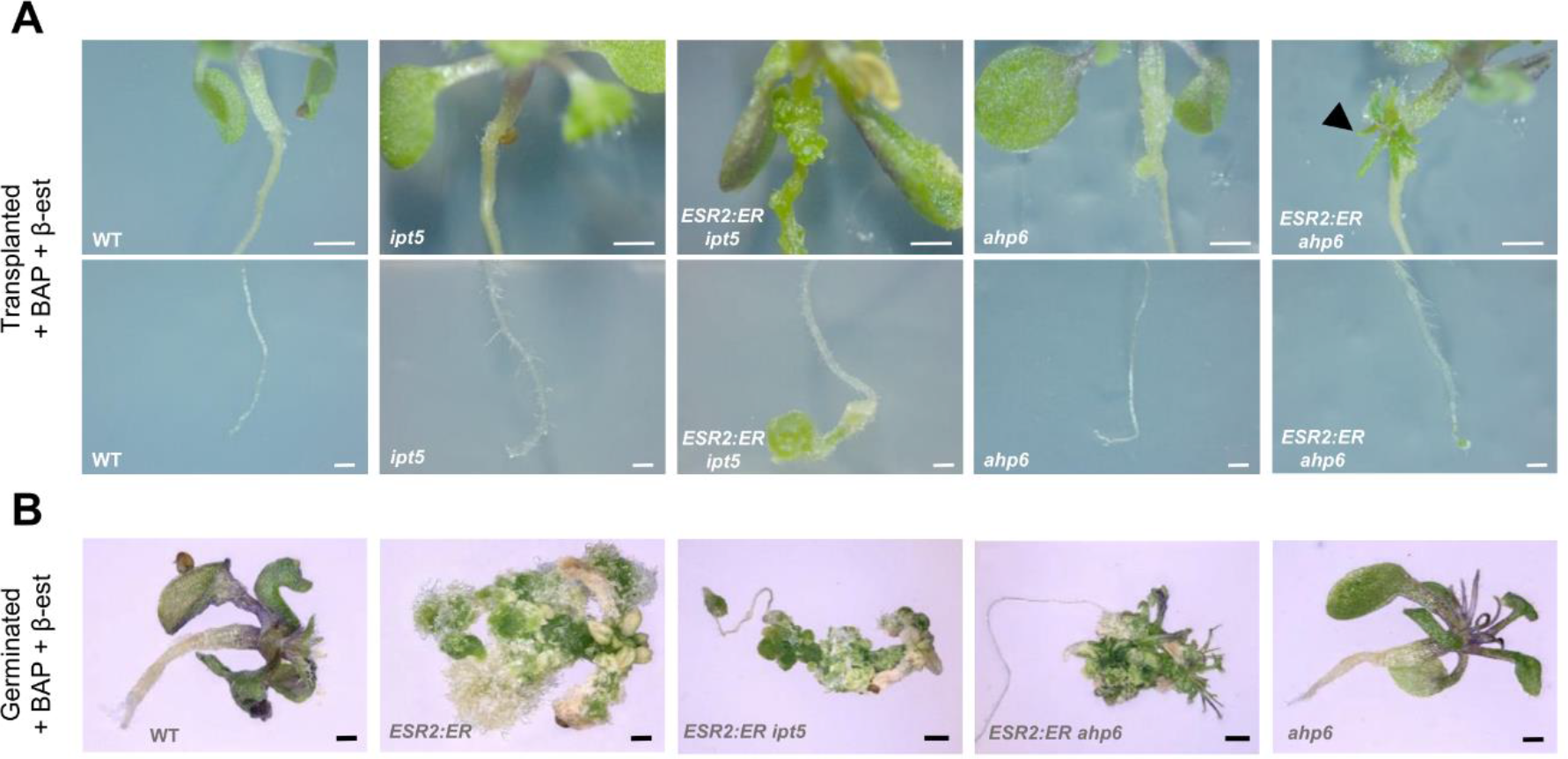
Effect of exogenous cytokinin application in calli development in *ESR2:ER ipt5* and *ESR2:ER ahp6* seedlings. A) Seedlings grown in medium supplemented with 20 µM of β- estradiol and transplanted at 6 DAG to media with 30 µM of 6-benzylaminopurine (BAP) and 20 µM of β-estradiol, observed after 34 days after transference. A black arrowhead indicates a shoot- like structure in the hypocotyl-root junction. **B)** Seedlings directly germinated in medium supplemented with 30 µM of 6-benzylaminopurine (BAP) and 20 µM of β-estradiol, observed at 40 DAG. Scale bars: A) shoots 500 µm and roots 200 µm; B) WT, *ESR2:ER,* and *ahp6* 500 µm, *ESR2:ER; ESR2:ER ahp6* and *ESR2:ER ipt5* 1 mm.

We next evaluated callus formation in *ESR2:ER, ESR2:ER ipt5,* and *ESR2:ER ahp6* seedlings that were directly germinated in a medium supplemented with BAP and β-estradiol. They were observed 40 days after induction. In these conditions, induced *ESR2:ER* plants became very compact, lacked visible roots, and the whole plant transformed largely into a large mass of calli (Fig. 4B).

Direct germination in these conditions intensified calli formation in *ESR2:ER ipt5* plants compared to those transferred to the BAP medium after germination. They exhibited extensive callus formation (Fig. 4B), resembling induced *ESR2:ER* plants to a greater extent. Calli developed in the root, only not along the entire root as in *ESR2:ER* plants (Fig. 4B). Instead, <u>c</u>alli developed mainly at the hypocotyl, its base, the root-hypocotyl junction, and the root apex. Again, though the formation of calli was still more pronounced in *ESR2:ER* than *ESR2:ER ipt5* plants, BAP addition to the medium promoted increased callus formation in *ESR2:ER ipt5* plants in comparison to the same plants grown in induction medium without this hormone.

*ESR2:ER ahp6* seedlings presented visible roots without calli. Instead, calli developed from aerial tissues (Fig. 4B). *ESR2:ER ahp6* shoots were severely disorganized. Interestingly, besides calli, shoot-like structures, leaves, and some radialized, rod-like structures could be recognized in the aerial part of these *ESR2:ER ahp6* plants (Fig. 4B). It is worth noting that calli did not develop in any of the control plants (WT, *ahp6*, and *ipt5*) under the tested conditions. In some BAP-treated *ahp6* seedlings, a slight growth of ectopic tissue was observed in the hypocotyl. This tissue was not green, and its growth was moderate in comparison to ESR2-induced calli that had developed at the same time as green cell masses (Fig. 4B).

In conclusion, these experiments indicate that cytokinin biosynthesis mediated by *IPT5* has a role in callus formation induced by ESR2. Increased cytokinin levels, on the other hand, partially restored callus formation in *ESR2:ER ahp6* roots to a lesser extent than in *ESR2:ER ipt5*.

### ESR2 can bind to *IPT5* and *AHP6* regulatory regions and activate their expression

After confirming that both *IPT5* and *AHP6* were upregulated and contributed to the phenotypic alterations caused by ESR2 overexpression, we explored the nature of the regulation of *IPT5* and *AHP6* by ESR2. The RT-qPCR experiments in seedlings treated with cycloheximide suggested a possible direct regulation; hence, to further investigate this hypothesis, we employed Yeast-1- Hybrid and NanoLUC transactivation assays. Many AP2/ERF-type transcription factors recognize a conserved binding site, the GCC box (Fujimoto et al., 2000; Hao et al., 1998; Ohme-Takagi & Shinshi, 1995). Previous works have shown that ESR2 binds to and regulates targets through a GCC box (Eklund et al., 2011; Zhang et al., 2018). Using the Plant Promoter Analysis Navigator (PlantPAN; http://PlantPAN.itps.ncku.edu.tw/ (Chow et al., 2019), we identified one GCC box at -289 bases from the *IPT5* ATG (pink diamonds in Fig. 5A). In addition, three other sequences related to the GCC box were also identified in *IPT5*, two of them within the *IPT5* ORF, 10 and 18 bases near the translation stop codon and another in the 3’ UTR (purple diamonds, Fig. 5A). We considered these regions as potential ESR2 binding sites and evaluated them using Y1H assays.

**Figure 5.**
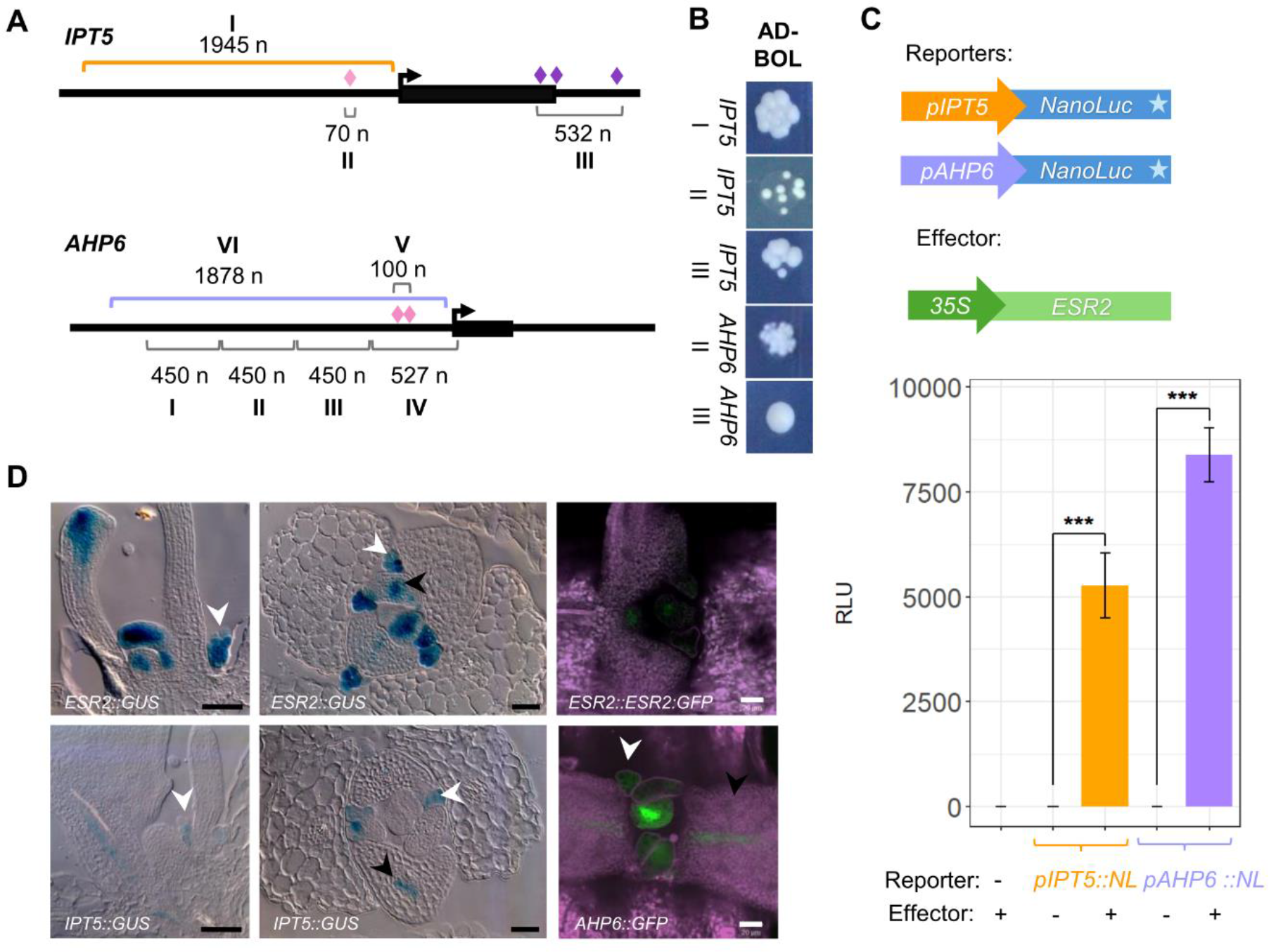
ESR2 regulation of *IPT5* and *AHP6* expression and correlation with ESR2 expression. A) Schemes of *IPT5* and *AHP6* genomic regions tested by Y1H (*IPT5* I to III, *AHP6* I to V fragments) and NanoLuc assays (purple and yellow regions). Diamonds indicate GCC sites before the ATG of each gene (pink) or after (purple). **B)** Y1H assays for ESR2 protein interaction with *IPT5* and *AHP6* fragments. **C)** NanoLuc assays, the diagrams show the constructs used as effector and reporters. Relative NanoLuc activity detected in each combination is presented in the graph. Values are given as mean ± SE. Significant differences are indicated by asterisks (***p <0.001 in Student’s t-test). **D)** *ESR2* expression compared to *IPT5* and *AHP6* expression in aerial tissues of seedlings. From right to left, the first two microphotographs show longitudinal sections of *ESR2::GUS* and *IPT5::GUS* seedlings. Microphotographs in the middle show transversal sections of *ESR2::GUS* and *IPT5::GUS* seedlings. The last microphotographs show *ESR2::ESR2:GFP* and *AHP6::GFP* expression. White arrowheads indicate stipules and black arrowheads indicate expression in the vascular tissue. Scale bars: 50 um in GUS histological sections and 20 um in GFP micrographs.

We tested two fragments comprising the region upstream from the ATG: A 1945 bp (fragment I) and a 70 bp long fragment that contained the GCC box (fragment II). In addition, a fragment of 532 bp that included the three GCC boxes located at the end of the *IPT5* ORF and its 3’ UTR (fragment III) was also tested. The results of the Y1H assays revealed an interaction between ESR2 and the 3 fragments (Fig. 5B).

In the case of the *AHP6* promoter, we identified two boxes at -258 and -293 bases from the start codon (pink diamonds) (Fig. 5A). Through the Y1H assay, we tested five *pAHP6* promoter fragments, including fragments I to III (450 bases each), fragment IV (527 bp), and V (a small fragment of 100 bp) containing the region of GCC boxes (Fig. 5A). However, fragments I, IV and V presented auto-activation (the activation of reporter genes without the participation of the transcription factor) and could not be tested. Promoter fragments II and III did not show autoactivation and the Y1H assays revealed ESR2 interaction with them (Fig. 5B).

We then validated the regulation of *IPT5* and *AHP6* by ESR2 *in planta* using a transient luciferase assay. We tested promoter fragments comprising 1945 bp upstream of the ATG of *IPT5*, and 1900 bp upstream of the ATG of *AHP6*. Both promoter fragments were fused to the bioluminescent reporter NanoLuc (NL). These constructs were transiently expressed in tobacco leaves together with a *35S::ESR2* construct. In the absence of the *35S::ESR2* construct, we did not detect NanoLuc activity (Fig. 5C). However, when *35S::ESR2* was co-infiltrated with *pIPT5::NL* or *pAHP6::NL,* a clear increase in bioluminescence was detected (Fig. 5C). The results of both the Y1H and NanoLuc experiments suggest direct binding and activation of *pIPT5* and *pAHP6* by ESR2.

Subsequently, to investigate whether the regulation of ESR2 towards *IPT5* and *AHP6* could occur in the natural context of *ESR2* expression, we compared their natural expression patterns during seedling development. We explored *IPT5* expression using the *IPT5::GUS* promoter reporter line and compared it with the *ESR2* promoter marker line *BOL::GUS* (Marsch-Martinez et al., 2006). Visualization of stained *IPT5::GUS* whole seedlings revealed overlapping in some tissues where *ESR2* is also expressed (Fig. S5). High overlapping between *ESR2* and *IPT5* expression could be mainly observed in young leaf tissues, particularly leaf vasculature at very early stages of development and stipules (Fig. 5D). In more developed tissues, diverse expression patterns were evident since *IPT5* was expressed in the vasculature of cotyledons and leaves at later stages of development, while the expression of *ESR2* remained confined to discrete points in the apex of young leaves and cotyledons (Fig. S5).

We also analyzed *AHP6* expression using the *AHP6::GFP* reporter line and compared it to a genomic reporter line, *ESR2::ESR2:GFP*. As previously reported, *ESR2* expression was observed in the meristem peripheral zone, at the region where a new leaf primordium will emerge, and in leaf primordia (Fig. 5D, Ikeda, et al., 2006a; Marsch-Martinez et al., 2006b; Nag et al., 2007). Moreover, *ESR2* expression was also observed in tissues such as the developing main vasculature of the youngest leaves and the stipules (Fig. 5D). *AHP6* expression coincided in leaf primordia, stipules, and the developing vasculature of young leaves (Fig. 5D). In summary, these results suggest that the regulatory effect of ESR2 upon *IPT5* and *AHP6* could be direct and that these genes might be natural targets in specific tissues of the plant, such as the stipules and young vasculature for *IPT5*, and these tissues plus leaf primordia for *AHP6*.

## Discussion

Our study aimed to determine whether the transcription factor ESR2 could modulate the cytokinin pathway positively. Callus formation plays a crucial role in indirect regeneration (reviewed by Ikeuchi et al., 2013, 2019) and can be triggered by specific ratios of the phytohormones cytokinin and auxin. Subsequently, shoot regeneration occurs under high cytokinin to low auxin ratios (Ikeuchi et al., 2019; Skoog & Miller, 1957).

ESR2 has a remarkable ability to induce green callus formation and shoot regeneration even without the addition of exogenous hormones (Ikeda et al., 2006a; Marsch-Martinez et al., 2006) and the effects of ESR2 induction are comparable to those observed with cytokinin treatments or increased cytokinin content in plants. Other lines of evidence further suggest a potential positive role of ESR2 in cytokinin-related processes. ESR2, together with REVOLUTA (REV), positively regulates *SHOOT MERISTEMLESS (STM)* (Zhang et al., 2018), and STM promotes cytokinin biosynthesis by activating the expression of *IPT7* (Jasinski et al., 2005; Yanai et al., 2005). Moreover, alterations in the effects of cytokinin treatments in young gynoecia of *drnl-2* (an ESR2 loss-of-function allele (Nag et al., 2007)) mutants indicated a positive role in the early developmental stages of this important reproductive organ (Durán-Medina, et al., 2017a). Furthermore, global expression analyses in *lfs* (loss of function of the only *ESR2* homolog in tomato) also supported a potential positive effect of *ESR2* on the cytokinin pathway (Capua & Eshed, 2017).

In line with these observations, when we measured different cytokinin forms in ESR2-induced plants, a clear increase in total cytokinins was observed, especially iP and cZ types. In particular, iP types have been associated with the ability of cytokinins to promote root primordia disorganization and shoot identity (Pernisova et al., 2018); these effects are also observed in ESR2 overexpressors. Isopentenyltransferases (IPTs) catalyze a critical step in cytokinin biosynthesis, and *ipt* mutants have reduced levels of iP and tZ types (Miyawaki et al., 2006), while *IPT* activation lines or overexpressors predominantly exhibit an increase in iP types (Kakimoto, 2001; Sun et al., 2003; Zubko et al., 2002). Increased *IPT* expression enhances shoot regeneration and leads to phenotypes such as green callus formation in cytokinin-free conditions, shorter roots, and dark green cotyledons (Kakimoto, 2001; Sun et al., 2003; Zubko et al., 2002). These phenotypic similarities with ESR2/BOL overexpression lines (Ikeda et al., 2006; Marsch-Martinez et al., 2006) and the higher iP-type cytokinin content pointed to IPTs as potential downstream genes responsible for the increased cytokinin content and related phenotypes associated with ESR2. In tissue culture, callus formation has been linked to a lateral root development program (Atta et al., 2009; Sugimoto et al., 2010). *IPT5*, among the potential candidate *IPTs*, exhibits increased expression during the early stages of both lateral root and callus formation (Atta et al., 2009), and showed upregulation in roots and shoots just 30 minutes after ESR2 induction. This initial response might have been overlooked in previous studies because the increase in expression was moderate at this short time and the ESR2 activity leakage in the inducible line. Nevertheless, *IPT5* expression gradually intensified over time, coinciding with the formation of calli triggered by ESR2 induction.

Intriguingly, while ESR2 induction caused increased *IPT5* expression and cytokinin content, *AHP6*, a negative regulator of cytokinin signaling (Mähönen et al., 2000), has also been found to be activated by ESR2 in other tissues. ESR2 activates *AHP6* in root explants (Ikeda et al., 2006); in young gynoecia and inflorescences (Durán-Medina et al., 2017a); and has been found as an ESR2 target in lateral organ founder cells (Frerichs et al., 2019). The results obtained in the Y1H and transient expression experiments further suggested that ESR2 binds and activates the promoters of *IPT5* and *AHP6* directly. Notably, *AHP6* has been recently found to be a direct target in inflorescence meristems (Dai et al., 2023).

Therefore, interestingly, ESR2 induction can activate both cytokinin biosynthesis (*IPT5)* and negative signaling (*AHP6*), and some ESR2-induction phenotypes, such as initial root growth inhibition, were differently affected by the loss of each gene. However, despite their seemingly contradictory effects in the cytokinin pathway, both functional *IPT5* and *AHP6* are essential to obtain other ESR2-induced phenotypes, such as stunted leaf formation. Remarkably, they are both required for the massive green calli formation in roots, one of the most conspicuous phenotypes caused by ESR2 activation. This suggests that ESR2-mediated callus induction requires a delicate balance between an increase in cytokinin content and a reduction in cytokinin signaling in the regions where *AHP6* expression is upregulated.

Cytokinins are known to move from cell to cell (reviewed by Durán-Medina et al., 2017b; Liu et al., 2019). Free CK bases, the biologically active forms, have been proposed to diffuse through cell membranes (Nedvěd et al., 2021). Building on this knowledge, we propose a scenario to explain the dual requirement for cytokinin biosynthesis and negative regulation during ESR2 - induced callus development, in which ESR2-responsive regions are established where cytokinins are produced (Fig. 6). These regions would serve as hubs for cytokinin synthesis, which is then mobilized through transport or diffusion to neighboring cells. Simultaneously, cytokinin signaling is selectively reduced within the ESR2-responsive region. However, this reduction does not extend to neighboring cells, which respond to the cytokinin produced in the adjacent regions, creating differential CK response fields. ESR2 would then modulate the delicate balance between cytokinin production and localized signaling modulation, which leads to callus formation in this system. Such positive-negative cytokinin regulation modules affect the other tissues of the plant. Positive regulation of both cytokinin biosynthesis and negative signaling or degradation is required for proper vascular development. The TARGET OF MONOPTEROS 5/LONELY HIGHWAY (TMO5/LHW) dimer activates cytokinin biosynthesis through *LONELY GUY 3* and *4 (LOG3* and *4*) as well as reduces cytokinin signaling through the activation of *AHP6* in xylem precursor cells (Ohashi-Ito et al., 2014). Moreover, the TMO5/LHW dimer also activates cytokinin degradation indirectly via *CKX3* expression in adjacent cells (Yang et al., 2021). A similar scenario could occur in the case of ESR2-induced calli (Fig. 6), which warrants investigating whether this is the case, in these and other calli. Curiously, *ESR2* has also been reported to be a target of MONOPTEROS (MP) (Dai et al., 2023).

**Figure 6.**
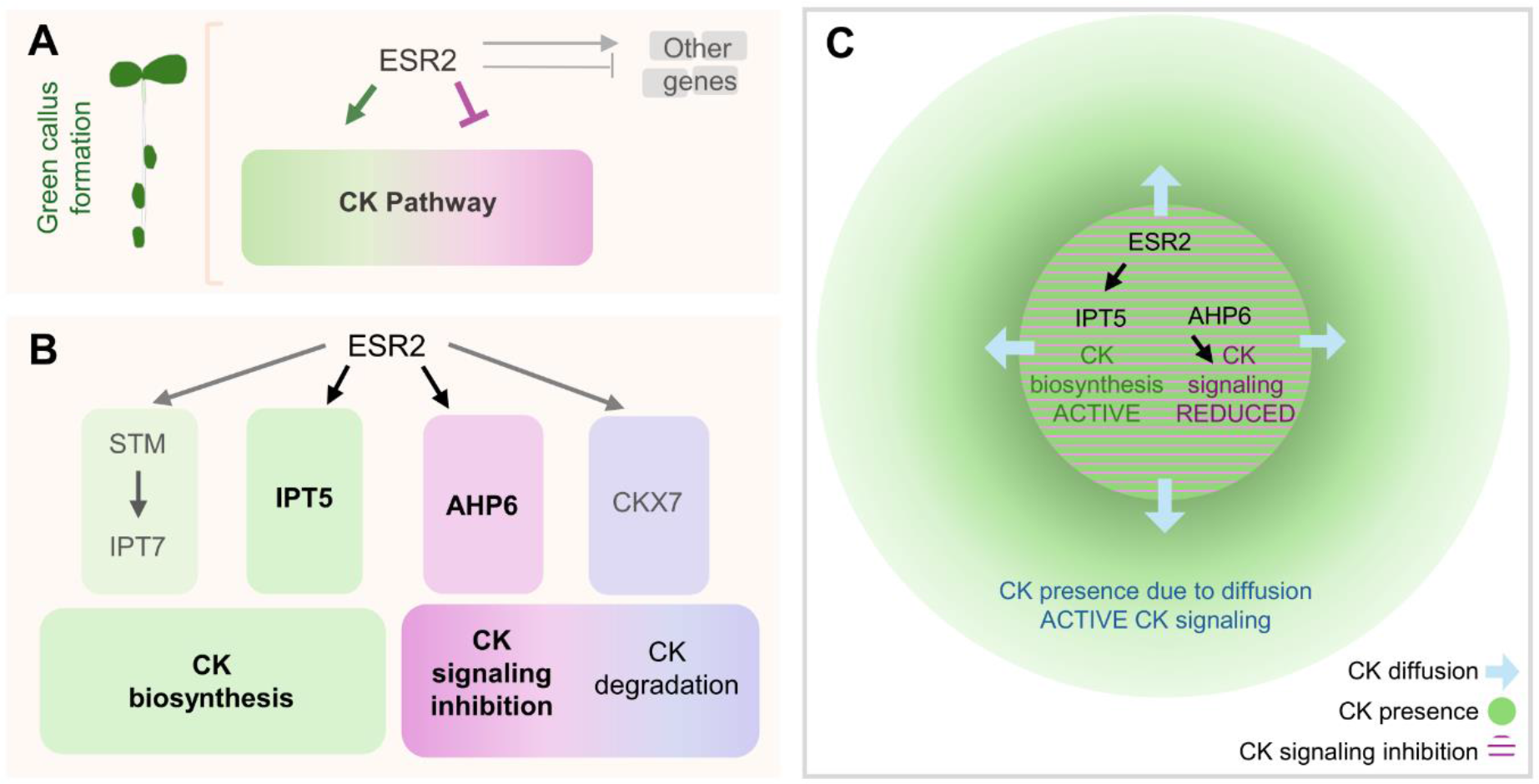
Model of ESR2 action in the cytokinin pathway leading to root reprogramming and green callus formation. A) Callus formation and other phenotypes promoted by the induction of ESR2 activity depend on its modulation of the cytokinin pathway, both positively and negatively. Other ESR2-regulated genes (cytokinin or other processes) are also involved. **B)** Cytokinin genes regulated by ESR2 and their effect in the cytokinin pathway. The genes described by Zhang et al 2018, Jasinski et al., 2005, Yanai et al., 2005, Dai et al, 2023, Frerichs et al. 2019; Ikeda et al., 2006 and in this work are depicted. The genes in bold were studied in this work, and when mutated, abolish or severely reduce ESR2-promoted callus formation. **C)** Proposed model of one possible scenario to explain the effect of ESR2 in cytokinin biosynthesis and inhibition of cytokinin signaling, creating regions where cytokinins are produced but signaling is reduced, and regions to where cytokinins diffuse and signaling is not inhibited. These differential cytokinin sensing “fields” could trigger the reprogramming of root cells to form green calli.

As expected, given the function of these genes, differences in callus formation were observed in *ESR2:ER ahp6* and *ESR2:ER ipt5* mutants when supplemented with cytokinins, since this ability was recovered to a greater extent in *ESR2:ER ipt5* than in *ESR2:ER ahp6* plants. Interestingly, in the induced *ESR2:ER ahp6,* supplemented with cytokinins, radialized aerial organs were formed, reminiscent of the ones observed in *drn-D*, the activation tagging line of one of the closest homologs to ESR2, ESR1/DRN, gain-of-function mutants (Kirch et al., 2003). Moreover, shoot formation in the hypocotyl-root junction was obtained in these lines, which was never observed before in *ESR2:ER* whole seedlings in the wild-type background, where only calli develop after induction. Therefore, *AHP6* could contribute to the inhibition of proper shoot regeneration from green calli in these seedlings.

Cytokinin treatments alone were not enough to trigger equivalent ESR2-induced callus formation in wild-type, non-induced *ESR2:ER*, *ipt5* or *ahp6* plants, indicating that the increase in cytokinin biosynthesis in these lines is not enough to induce calli, and that ESR2 regulates other processes in combination with cytokinin biosynthesis and negative signaling, which are also required for callus development. One of these processes could be related to auxin (Chandler et al., 2011; Chandler & Werr, 2014; Skoog & Miller, 1957). ESR2 activates *SHORT INTERNODES/STYLISH* (*SHI/STY*) (Eklund et al., 2011), which in turn activates auxin biosynthesis (Sohlberg et al., 2006). Nevertheless, future studies should identify the other genes (including other auxin or cytokinin-related genes) and processes that are involved in ESR2-induced callus formation.

Interestingly, while the induced activation of ESR2, whose expression is driven by a 35S promoter, was expected to upregulate *IPT5* and *AHP6* indistinctly in many plant tissues, the upregulation of both genes was mostly restricted to the vascular region (in the analyzed roots and shoots). This might indicate that other factors, including epigenetic marks or interacting transcription factors, among others, restrict the action of ESR2 on these genes and confine it to the vascular tissues. Moreover, the induced plants also presented an enlarged vascular region (Fig. 3D). Cytokinins promote procambial cell proliferation and secondary growth (Matsumoto-Kitano et al., 2008; Ye et al., 2021). The increased number of vascular cells observed after ESR2 induction was reduced in the *ipt5* mutant, suggesting that it could be due to an increase in cytokinin biosynthesis. Conversely, the lack of *AHP6* function exacerbated the phenotype, observed as a larger region of lignified cells, possibly due to an increased sensitivity to the high cytokinin content.

### Natural context and relevance of regulation

We studied the regulation of the *IPT5* and *AHP6* cytokinin genes by ESR2 in the context of its ability to induce callus formation and other cytokinin-related phenotypes when overexpressed, and it is possible that this regulation could occur in a natural context. Co-expression can provide insights into where this regulation could occur naturally.

Expression of both *ESR2* and *AHP6* has been reported in founder cells of floral organs (Besnard et al., 2014; Chandler et al., 2011), where the regulation could occur naturally, and, accordingly, ESR2 has been confirmed to activate *AHP6* in the aerial shoot meristem (Dai et al., 2023).

Interestingly, we observed co-expression of *ESR2*, *AHP6*, and *IPT5* in the often-overlooked stipuli within the developing primordia of the vegetative shoot apical meristem. Their function remains largely unknown, but they have been implicated as sites of diffusible signal production (Oppenheimer et al., 1991). It would be interesting to further investigate whether the expression of these three genes indicates that stipuli could be an insensitive reservoir of cytokinins, capable of diffusing to the developing organs or even the SAM.

Another region reflecting the coincidence of the three genes was the vasculature of developing leaves. Coupled with the sharp increase in p*IPT5* and p*AHP6* expression upon ESR2 induction in the vascular region, this observation hints at ESR2’s potential role in naturally activating these genes in the specific tissue. Interestingly, ESR2 was found, and recently confirmed to directly activate *CKX7* (Dai et al., 2023; Ikeda et al., 2006), which is also expressed in the vasculature (Köllmer et al., 2014). We could speculate that ESR2 may also have a role in promoting the formation of differential cytokinin content or sensibility fields during vascular development, by inducing cytokinin biosynthesis and promoting insensibility or degradation in the cells where it is expressed. Interestingly, a previously unidentified role for ESR2/DRNL in vascular development has been recently uncovered, supporting this hypothesis (Glowa et al., 2021), and the increased number of vascular cells after ESR2 induction also suggests ESR2’s role in this process. Nevertheless, it is also possible that the regulation of *IPT5, AHP6*, and *CKX7* by ESR2 occurs at different stages or tissues (for example, during flower or floral organ development), and this should be explored in further detail in other investigations.

**In summary,** ESR2 influences both cytokinin levels and negative cytokinin signaling that lead to callus formation and other phenotypes, and these effects may be ESR2’s natural function in guiding plant development.

## MATERIALS AND METHODS

### Plant materials and growth conditions

The lines used in this study were: Columbia (Col-0) wild-type plants; *ipt5-2* (Miyawaki et al., 2006) and *ahp6-1* mutants (Mähönen et al., 2006, genotyped as in Muller et al., 2017); *pBOL::BOL:GFP* (this work, renamed as *ESR2::ESR2:GFP*), *BOL::GUS* (Marsch-Martínez et al., 2006), *IPT5::GUS* (Dello Ioio et al., 2008), and *AHP6*::*GFP* (Mähönen et al., 2006); and the inducible ESR2 line *35S::ESR2-ER* (Eklund et al., 2011; Ikeda et al., 2006), mentioned in this work as *ESR2:ER.* More information in the Supplementary Material and methods section.

### ESR2::ESR2:GFP reporter construct and line

A 6 kb fragment containing regulatory sequences upstream the ATG of *DRNL/ESR2/BOL* and the coding region until just before the stop codon was amplified from wild type Col-0 DNA, using Phusion polymerase (New England Biolabs) and cloned in the pENTR/D-TOPO vector, using the corresponding cloning kit (Invitrogen). The fragment was confirmed by sequencing, and the promoter fragment was recombined using Gateway LR Clonase Enzyme mix (Invitrogen) into the pMDC204 vector (Curtis & Grossniklaus, 2003), resulting in construct *ESR2::ESR2:GFP*. The resulting binary vector was transformed to *A. tumefaciens* pGV2206 and, using a modified floral dip method, introduced in *drnl-2* (Nag et al., 2007) plants. Transformants were selected in hygromycin-containing medium.

### Histology and microscopy

*ESR2:ER* induction was either performed in 6 DAG seedlings germinated in 0.5x MS medium and transferred to medium supplemented with 10 μM β-estradiol, or germinated directly in the latter medium, and compared to those transplanted to or germinated in mock MS medium supplemented only with DMSO (the solvent used to dissolve β-estradiol). To determine the regulation of *AHP6* and *IPT5* expression in response to ESR2 induction in specific tissues, crosses between *ESR2:ER* and *AHP6::GFP* or *IPT5::GUS* expression marker lines were obtained. Homozygous *ESR2:ER AHP6::GFP* and *ESR2:ER IPT5::GUS* seedlings were transplanted to MS medium with 10 μM β- estradiol (Sigma–Aldrich). Treated seedlings were removed from the plates and processed for expression visualization 48 hours after ESR2 induction. More information in the Supplementary methods section.

### Gene expression analysis by RT-qPCR

For RT-qPCR analysis, nine days old (DAG) Columbia and *ESR2:ER* seedlings were sprayed with different solutions: (a) 10 μM β-estradiol and 30 μM cycloheximide (CHX), (b) 30 μM cycloheximide, and (c) DMSO (solvent) in distilled water. Solution (a) was the inductor solution, β-estradiol promotes nuclear transport of the chimeric ESR2:ER transcription factor, while solution (b) and (c) were used as mock. Cycloheximide was used to prevent new protein synthesis. Aerial tissue was collected 30 minutes after treatment application. Table S1 includes the sequences of the primers used. More information in the Supplementary Material and methods section.

### Calli development evaluation and cytokinin treatments

To analyze the relevance of *AHP6* and *IPT5* in ESR2-promoted callus development, crosses were made between *ESR2:ER* and the *ahp6-1* and *ipt5-2* mutants. The *ahp6-1* allele was genotyped according to (Müller et al., 2017) and *ipt5-2* was genotyped by PCR using gene and T-DNA specific primers. For the evaluation of callus development, Col WT, *ESR2:ER,* single mutant alleles*, ESR2:ER ahp6* and *ESR2:ER ipt5* seedlings were grown in MS 0.5x medium with 20 μM β-estradiol and callus development was evaluated after 40 days. To test the effect of cytokinin in the recovery of callus formation, the same seedlings were grown in MS 0.5x medium with 20 μM β-estradiol and at 6 DAG were transplanted to medium with 20 µM of β-estradiol and 30 µM of 6-benzylaminopurine (BAP, Duchefa Biochemie) or medium with 20 µM of β-estradiol and the mock solution NaOH (0.2 N, which was the solvent used for BAP), or were germinated directly in these media. More information in the Supplementary Material and methods section.

### Promoter analyses

*In silico* analyses of the *AHP6* and *IPT5* regulatory sequences were made with the Plant Promoter Analysis Navigator (PlantPAN; http://plantpan.itps.ncku.edu.tw/plantpan3/, (Chow et al., 2019)) to identify putative GCC box-like regulatory element sequences.

### Y1H assays

The binding of ESR2 to the promoter regions of putative target genes was tested using a Yeast one-hybrid system (Y1H), based on the Matchmaker Gold Yeast One-Hybrid protocol (https://www.takarabio.com). Promoter regions of *IPT5* and *AHP6* (Fig. 5A) were amplified using primers indicated in Table S1. See Supplementary Material and methods.

### NanoLuc activity assays

Promoters of *IPT5* and *AHP6* were recombined into the binary vector pGWB601-NL3F10H (Urquiza-García & Millar, 2019) using LR clonase mix (Invitrogen), resulting in the *pAHP6::*NL and *pIPT5::*NL reporter constructs, which were transformed into *Agrobacterium tumefaciens* GV3101. Transient assays were performed as described in (Becerra-García et al., 2023). See Supplementary Material and methods.

### Quantification of endogenous cytokinins

Quantification of CK metabolites was performed as previously described in (Svačinová et al., 2012). Seeds of Col-0 and *ESR2:ER* were stratified in MS medium (Murashige & Skoog, 1962) and germinated in petri dishes in a growth chamber (22°C and 16 hours-light). Five days after germination, seedlings were transferred to MS with 10 µM β-estradiol (treatment medium) and MS without β-estradiol (“Mock” medium) and grown in the same conditions as before. Two weeks after the transference, around 20 mg of the treated and mock seedlings were harvested. They then were frozen with liquid nitrogen and stored at -80°C; Four independent biological replicates were harvested for each genotype and treatment. Samples (20-30 mgFW) were extracted in 1.0 mL modified Bieleski buffer (60% MeOH, 10% HCOOH and 30% H2O) together with a cocktail of stable isotope-labeled internal standards (0.25 pmol CK bases, ribosides, N-glucosides, and 0.5 pmol CK O-glucosides and nucleotides were added per sample). The extracts were purified using multi-StageTips (containing C18/SDB-RPSS/Cation-SR layers), the eluates were then evaporated to dryness in a vacuum and stored at -20°C. Cytokinin levels were determined using ultra-high performance liquid chromatography-electrospray tandem mass spectrometry (UHPLC-MS/MS) using stable isotope-labeled internal standards as a reference. Statistical analysis was performed using the Student’s t-test (*p < 0.05, **p < 0.01, ***p < 0.001).

## Acknowledgements

The authors thank Juan Carlos Ochoa Sánchez, Miriam Salvador Daniel, Elizabeth Mendoza Bravo, Francesco Florio, J. Irepan Reyes-Olalde, Joanna Serwatowska, Rosa Esmeralda Becerra, Juan Ramos and Vincent Cerbantez Bueno for technical help and material contribution. We are also grateful to Thomas Jack (*drnl-1* seed), Eva Sundberg (*ESR2:ER* seed), and Raffaelle Dello Ioio (for *IPT5::GUS* seed). The NMM Laboratory acknowledges the funding obtained through the CONAHCyT CB-G-2023-219 project, postdoc fellowships to Yolanda Durán and David Díaz, and Ph.D. fellowship to Herenia Guerrero (1008711). The SdF laboratory was financed by the CONAHCYT grants FC-2015-2/1061 and CB-2017-2018-A1-S-10126. SdF is grateful for the Fellowship of the Marcos Moshinsky Foundation (2018). NMM and SdF participate in the European Union project H2020-MSCA-RISE-2020 EVOfruland (101007738).

## Author contributions

Conceptualization, YDM, NMM; Methodology, YDM, HHU, JECV, LC, ON, SdF, NMM; Validation, YDM, DDR, MDM, AGF, JECV, HGL, ON; Formal analysis, DDR, MDM, JECV, HGL; Investigation, YDM, DDR, HHU, MDM, JECV, CAV, ON; Writing – original draft, YDM, NMM; Writing – review and editing, YDM, NMM; Visualization, YDM, DDR, HGL, NMM; Supervision, LC, ON, SdF, NMM; Project administration, NMM; Funding acquisition, NMM, LC, ON, SdF.

The authors declare no conflicts of interest.

## SUPPLEMENTARY MATERIAL

Supplementary Materials and Methods

Figure S1. IPT5 and AHP6 expression regulation by ESR2.

Figure S2. IPT5 expression during lateral root development.

Figure S3. Comparison of root length and development in ESR2:ER, ESR:ER ahp6 and ESR2:ER ipt5 seedlings after ESR2:ER activation.

Figure S4. Partial recovery of the ESR2 activation phenotype after 34 days in media with BAP and β-estradiol.

Figure S5. Comparison of ESR2 and IPT5 expression in Arabidopsis seedlings.

Table S1. Primers used in this work.

