## Supplemental figures for "ESR2 orchestrates cytokinin dynamics leading to developmental reprogramming and green callus formation"

Supplementary figures

Duran-Medina et al.,

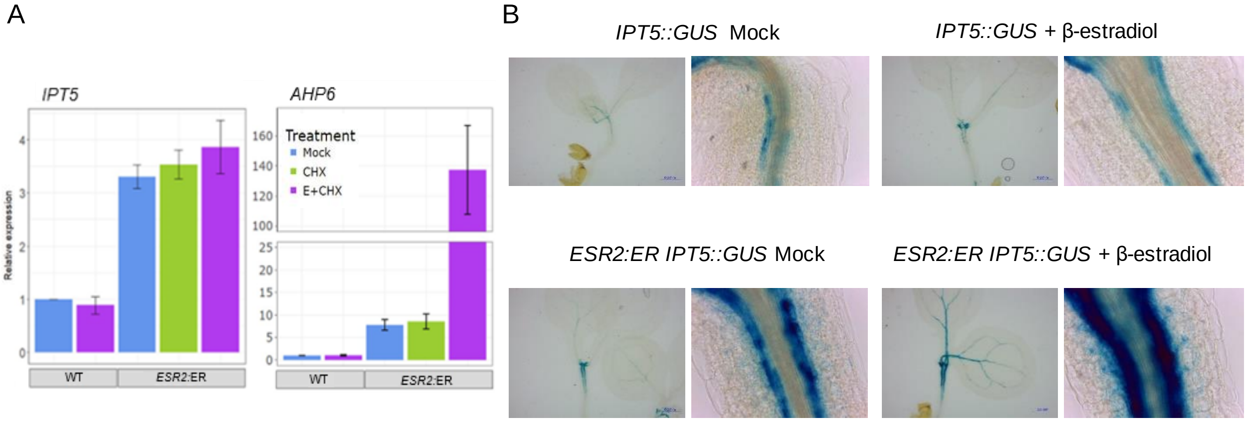


**Figure S1. *IPT5* and *AHP6* expression regulation by ESR2. A)** RT-qPCR analyses of *AHP6* and *IPT5* expression evaluated 30 minutes after ESR2 induction. Error bars indicate the S.E. **B)** *IPT5* expression in aerial tissues 48 hours after ESR2 induction, compared to the marker line in the wild type background.

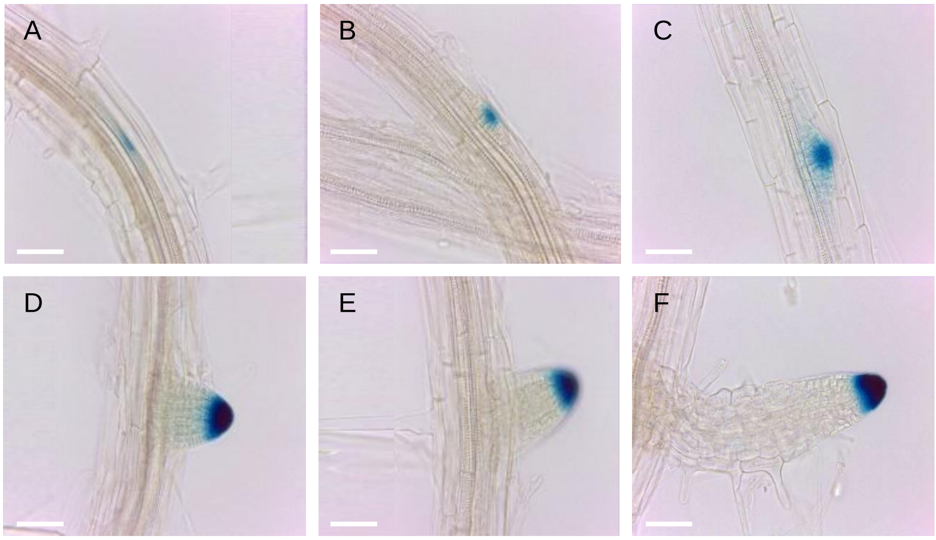


**Figure S2.** ***IPT5* expression during lateral root development**. **A) to C)** Initial stages of lateral root development. **D) to F)** Later stages of lateral root development. **A)** *IPT5* expression in adjacent cells to the xylem. **B)** Initial cell divisions. **C) to F)** *IPT5* expression maintenance at the root apex. Bars 0.05 mm

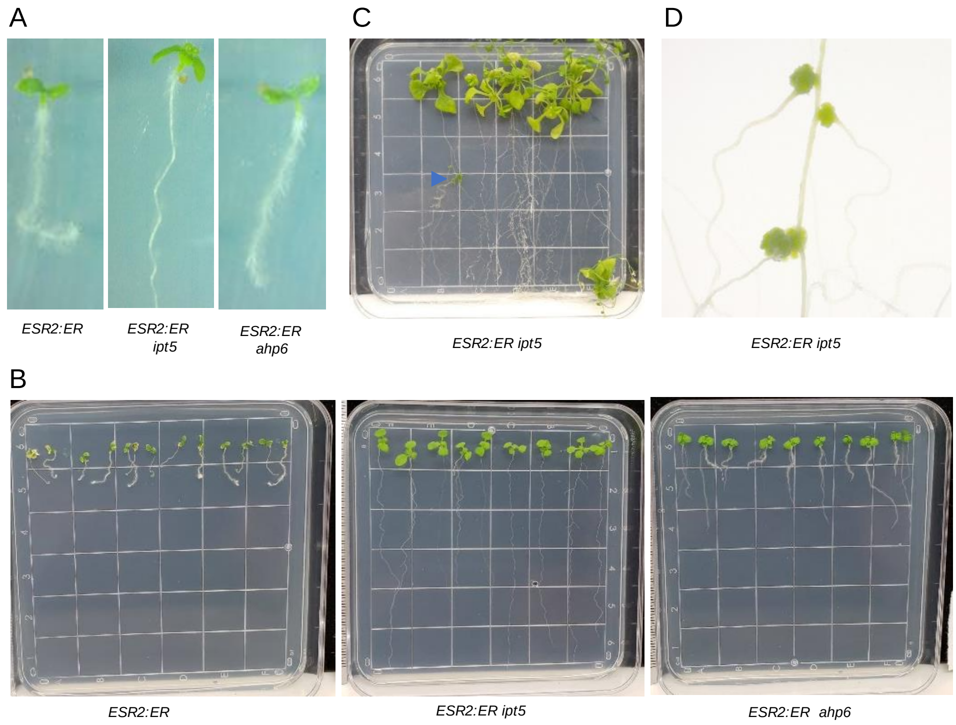


**Figure S3. Comparison of root length and development in *ESR2:ER*, *ESR2:ER ahp6* and *ESR2:ER ipt5* seedlings after *ESR2:*ER activation**. **A)** At 8 DAG. **B)** At 2 weeks after germination. **C)** *ESR2:ER ipt5* plants 40 days after ESR2 activation, little callus development (blue arrowhead). **D)** Magnification of the rare calli that developed at the base of lateral roots, in a few *ESR2:ER ipt5* plants, 40 days after ESR2 induction.

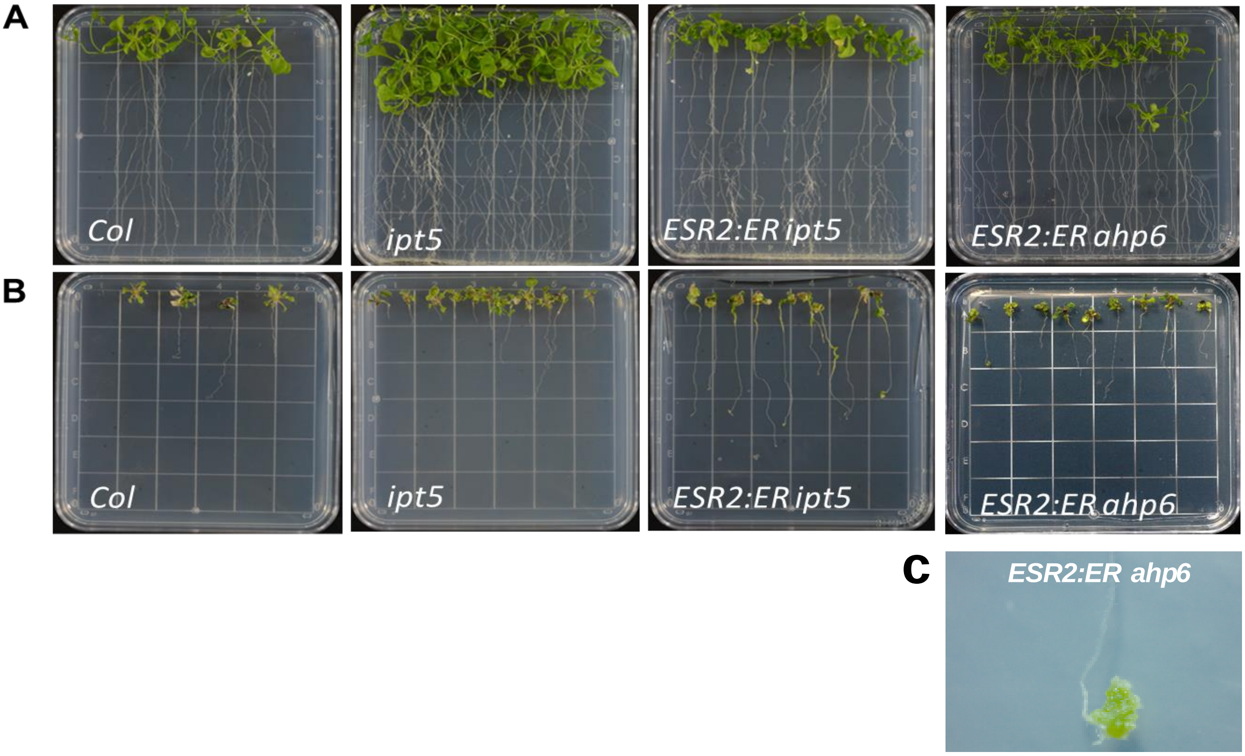


**Figure S4. Partial recovery of the ESR2 activation phenotype after 34 days in media with BAP and β-estradiol. A)** *S*eedlings grown in 20 µM β-estradiol medium and transplanted at 6 DAG to new 20 µM of β-estradiol medium (mock for the cytokinin treatment). **B)** *S*eedlings grown in 20 µM β-estradiol medium and transplanted to new 20 µM of β-estradiol medium supplemented with 30 µM 6-benzylaminopurine (BAP). **C)** Callus developed in the main root apex of a single *ESR2:ER ahp6* plant with β-estradiol and BAP.

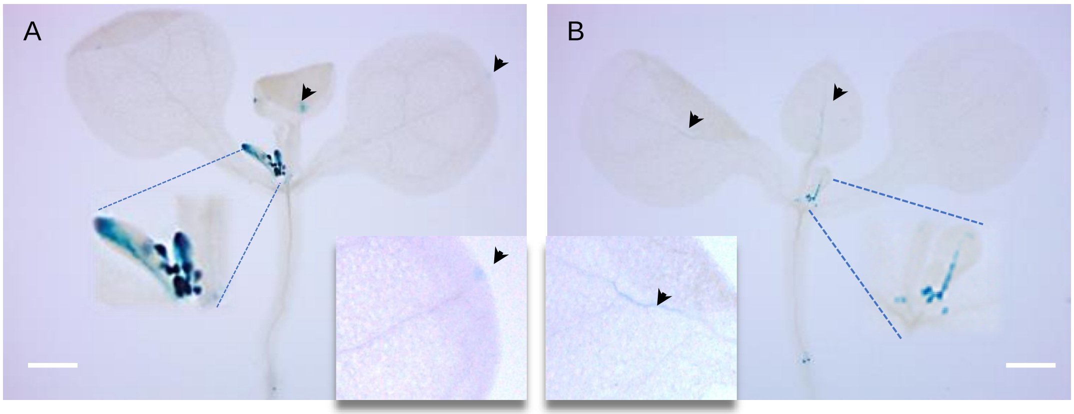


**Figure S5. Comparison of *ESR2* and *IPT5* expression in Arabidopsis seedlings**. **A)** *ESR2:GUS* expression. **B)** *IPT5:GUS* expression. Coincidences in expression are observed in some tissues, such as the young vasculature and stipules. Differences in mature cotyledons and young leaves at are highlighted by black arrowheads. Bars 500µm.
