## Supplemental Materials and Methods for "ESR2 orchestrates cytokinin dynamics leading to developmental reprogramming and green callus formation"

Durán-Medina et al.,

### Plant materials and growth conditions

All plants were grown *in vitro*. The seeds were disinfected with ethanol (70%) and sodium hypochlorite (20%), remaining 5 min in each solution. Subsequently, they were washed three times with distilled water, sown in MS 0.5x medium (PhytoTechnology Laboratories TM), 1.4% plant agar (PhytoTechnology Laboratories TM) and 1% sucrose, placed at 4 ° C for 48 hours; and finally transferred to a growth chamber for germination and growth, under long day conditions (16 h light and 8 h dark) at 22 ° C.

### Histology and microscopy

β-Glucuronidase staining and histological sections were described in (Durán-Medina et al., 2017). GUS-stained seedlings were observed under a Stemi 2000-C stereoscope coupled with Axiocam ERc 5s camera (Carl Zeiss) and a DM750 microscope coupled with ICC50 HD camera (Leica). GFP was observed using an LSM 510 META confocal scanning laser inverted microscope (Carl Zeiss). Propidium Iodide (PI; at 0.01 mg/mL) was used as a counterstain.

For xylem visualization, the *ESR2:ER*, *ESR2:ER ahp6*, and *ESR2:ER ipt5* lines were grown in MS medium with 20 μM β-estradiol (Sigma–Aldrich) and collected after 15 days. Hypocotyls and roots were collected and stained overnight using 0.2 % basic fuchsin (Sigma, cat no. 215597; St Louis MO, USA) in ClearSee (Kurihara et al. 2015; Ursache et al. 2018). For image acquisition, we used an LSM800 (Zeiss, Wetzlar, Germany) using the pre-loaded configuration for basic fuchsin visualization and a z-stack configuration. For the final pictures, maximum intensity projections were generated using 10 images.

### Gene expression analysis by qRT-PCR

Total RNA was extracted using the Quick-RNA™ MiniPrep kit (Zymo Research). The samples were treated with DNase I, included in the kit. Reverse transcription and amplification were performed using a KAPA SYBR FAST One-Step qRT-PCR Kit (Sigma-Aldrich) in a StepOne™

thermocycler (Applied Biosystems). Three technical and biological replicates were included in the analysis. Target gene expression levels were normalized to *ACTIN 2*. Data was analyzed using the $2^{-\Delta\Delta CT}$ method (Livak and Schmittgen 2001).

**Root measurements**

6-day old (DAG) *ESR2:ER* seedlings grown in MS 0.5x medium were transplanted to induction or mock medium. The plates were photographed 15 days after ESR2 induction and the pictures were analyzed using the Image J software (Schneider et al. 2012). Primary root length was measured, and the number of visible lateral roots was counted. In the case of root length analysis in the *ipt5-2* and *ahp6-1* backgrounds, the seedlings were germinated directly in mock or induction medium, the photographs were taken 13days after ESR2 induction and only the primary root was measured. Statistical analyses were performed with Statgraphics.

**Y1H assays**

Promoter regions of *IPT5* and *AHP6* (Fig. 5A) were amplified using primers indicated in Table S1, cloned in an Entry vector (pENTR-D topo, Invitrogen), and recombined with CZN1810, a Gateway-compatible version of pAbAi (Clontech; (Danisman et al. 2012). The vectors were linearized (*Bsp*119I, Thermo Scientific) and inserted into PJ69-α haploid yeast genome via homologous recombination to generate bait clones. Positive insertions were selected on SD GLUC medium lacking uracil (Sigma-Aldrich). To test intrinsic resistance to Aureobasidin A (AbA, Takara), the bait clones were plated on SD GLUC medium lacking uracil supplemented with increasing concentrations of AbA in the range of 0 to 700 ng $mL^{-1}$. ESR2 as prey clone (AD-BOL, pDEST22, Invitrogen; (Lozano-Sotomayor et al. 2016) in the haploid PJ69-A yeast was mated with the bait clones and grown in YPAD medium. Diploid yeasts (promoter + AD-BOL) were selected on SD GLUC medium lacking tryptophan and uracil. To test protein-DNA interactions, diploid yeasts representing the possible combinations were plated on SD GLUC medium lacking tryptophan and uracil, with different concentrations of AbA, 250 ng $mL^{-1}$ for *IPT5* fragments; 300 ng $mL^{-1}$ for *AHP6A*; 700 ng $mL^{-1}$ for AHP6. Yeast growth was scored on the sixth day of incubation at 30°C

**Assays of NanoLuc activity in plant extracts**

The luminescence signal (relative light units, RLU) was taken using a luminometer (LmaxII$^{384}$, Molecular Devices), with a 10 s integration time. Three luminescence measurement readings were taken every 15 min, and four biological replicates were performed in each assay. Each assay was repeated at least 3 times. Background controls were obtained by using leaf disks with no infiltration or infiltrated only with the empty reporter construct. A Student's t-test was used to determine the significance of relative NanoLuc activity differences.

**Danisman S, van der Wal F, Dhondt S, Waites R, de Folter S, Bimbo A, van Dijk AJ, Muino JM, Cutri L, Dornelas MC, et al.** Arabidopsis Class I and Class II TCP Transcription Factors Regulate Jasmonic Acid Metabolism and Leaf Development Antagonistically. Plant Physiol. 2012:**159**(4):1511. https://doi.org/10.1104/PP.112.200303

**Durán-Medina Y, Serwatowska J, Reyes-Olalde JI, De Folter S, and Marsch-Martínez N**. The AP2/ERF transcription factor DRNL modulates gynoecium development and affects its response to Cytokinin. Front Plant Sci. 2017:**8**:298950. https://doi.org/10.3389/FPLS.2017.01841/BIBTEX

**Kurihara D, Mizuta Y, Sato Y, and Higashiyama T**. ClearSee: a rapid optical clearing reagent for whole-plant fluorescence imaging. Development. 2015:**142**(23):4168–4179. https://doi.org/10.1242/DEV.127613

**Livak KJ and Schmittgen TD**. Analysis of Relative Gene Expression Data Using Real-Time Quantitative PCR and the 2−ΔΔCT Method. Methods. 2001:**25**(4):402–408. https://doi.org/10.1006/METH.2001.1262

**Lozano-Sotomayor P, Chávez Montes RA, Silvestre-Vañó M, Herrera-Ubaldo H, Greco R, Pablo-Villa J, Galliani BM, Diaz-Ramirez D, Weemen M, Boutilier K, et al.** Altered expression of the bZIP transcription factor DRINK ME affects growth and reproductive development in Arabidopsis thaliana. Plant J. 2016:**88**(3):437–451. https://doi.org/10.1111/TPJ.13264

**Schneider CA, Rasband WS, and Eliceiri KW**. NIH Image to ImageJ: 25 years of image analysis. Nature Methods 2012 9:7. 2012:**9**(7):671–675. https://doi.org/10.1038/nmeth.2089

**Ursache R, Andersen TG, Marhavý P, and Geldner N**. A protocol for combining fluorescent proteins with histological stains for diverse cell wall components. Plant J. 2018:**93**(2):399–412. https://doi.org/10.1111/TPJ.13784
