## Supplemental Table 1 for "ESR2 orchestrates cytokinin dynamics leading to developmental reprogramming and green callus formation"

| Name | Sequence | Description | Reference |
|---|---|---|---|
| ipt5 LP | CCTTTCCTCAACATATGCTCG | ipt5-2 genotipification | |
| ipt5 RP | CCACTGCTTGTAAAGCCTTTG | ipt5-2 genotipification | |
| AHP6-M FW | GAC GAG CAG TTC TTG CAG CTT CTG | ahp6-1 genotipification | Bartrina et al., 2010 |
| AHP6-M RV | CGT CAC GAA CCC TAC GAG CAC C | ahp6-1 genotipification | Bartrina et al., 2010 |
| IPT5 FW | CGATGACGAAAGAAGGGAAG | qRT-PCR | |
| IPT5 RV | CTCCAAGACAGCGACCAATC | qRT-PCR | |
| AHP6 FW | TAACGTCTGCGTTGCCTTT | qRT-PCR | Bishop et al., 2011 |
| AHP6 RV | CCTCCAGTCCTCTCAAGCAC | qRT-PCR | Bishop et al., 2011 |
| IPT5 I FW | CCGACTCCGATCAAACATGC | Y1H/NanoLUC (1945n) | |
| IPT5 I RV | TCGAGCTCTGGAACTCCAAT | Y1H/NanoLUC (1945n) | |
| IPT5 II FW | CACCcagtaacattcctccagcc | Y1H (70n) | |
| IPT5 II RV | GCTCTCAACGAGCATATG | Y1H (70n) | |
| IPT5 III FW | CACCCTG CCG TTC CGC CTC TTC | Y1H/NanoLUC | |
| IPT5 III RV | CAAACACGTATCGGACGGC | Y1H/NanoLUC | |
| AHP6 I FW | CACCATCTCAATGACTCATCATATC | Y1H | |
| AHP6 I RV | TACTTTTTCATGTTATCAGAGTAAGTAGT | Y1H | |
| AHP6 II FW | CACCTATGTTTAATACTAGCTAGTTG | Y1H | |
| AHP6 II RV | GAATAAAGTAAAAAATTATCAATAGTTAA | Y1H | |
| AHP6 III FW | CACCGTGAAACAATTTACG | Y1H | |
| AHP6 III RV | TTAATTTCTCACTGTTGAAATTAATCTC | Y1H | |
| AHP6 IV FW | CACCAGATAGAAAGTACATATTATTGGTG | Y1H | |
| AHP6 IV RV | CCACACCCAACCCCAACA | Y1H | |
| AHP6 V FW | CACCgctggtctgacagggtac | Y1H (100 n) | |
| AHP6 V RV | CACTCTGTGACCGGCTAC | Y1H (100 n) | |
| AHP6 VI FW | CATCTCAATGACTCATCATATCGAATGT | NanoLUC | |
| AHP6 VI RV | CCACAACGGCACACCCGT | NanoLUC | |
